# Mitosis sets nuclear homeostasis of cancer cells under confinement

**DOI:** 10.1101/2023.05.11.540326

**Authors:** Malèke Mouelhi, Alexis Saffon, Morgane Roinard, Hélène Delanoë-Ayari, Sylvain Monnier, Charlotte Rivière

## Abstract

During their life, mammalian cells are subjected to numerous mechanical constraints, especially in pathological contexts such as cancer. Recent studies have highlighted the central role of the nucleus in sensing mechanical cues, but they only focus on short periods of time, and so far, whether cells can adapt to prolonged confinement remains unknown. Here, we reveal the unsuspected role of mitosis in the long-term adaptation of nuclei to prolonged uniaxial confinement. For the colorectal cancer cell line investigated, following the first confined cell division, a new homeostatic state was reached by nuclei: they were smaller, and had reset the tension of their envelope. This adaptation through mitosis relied both on the nuclear tension sensor cPLA2 and the contractility machinery. We report for the first time a mechano-adaptation during mitosis, a process that could be crucial to adapt to stresses in the tumor microenvironment. We therefore anticipate that our work could provide new insight into cancer cell plasticity and cancer relapse.

**Significance Statement:** Most cell types undergo significant deformation throughout their life cycles. Immune cells must deform to navigate through dense matrices, while cancer cells in solid tumors experience squeezing from neighboring cells. The nucleus, central for many cell function, is the stiffest and largest organelle. Understanding its long-term response to spatial constraints is hence crucial yet largely unexplored.

In this study, we investigate how a colorectal cancer cell line adapts to prolonged confined environments, with a particular focus on nuclear dynamics under continuous squeezing.

Our groundbreaking findings reveal for the first time a mechano-adaptation during mitosis leading to a decrease in nuclear size.

This research contributes to the fundamental understanding of cellular mechanosensing, opening new avenues for cancer biology research.

## Introduction

The response of cancer cells to modifications in their microenvironmental mechanics has drawn attention over the last decades, since pioneering studies have shown that mechanics play an important role in the malignant transformation of cells during tumor progression and dissemination (1, 2). It is now acknowledged that changes in mechanical properties of the tumor microenvironment (densification and stiffening of the extracellular matrix, mechanical compression) foster a malignant and invasive phenotype (3) together with a decrease in cell proliferation (4, 5), and changes in cancer cell gene expression (6, 7). Hence, alleviating mechanical stress is now envisioned as an interesting therapeutic strategy (8–10). To better target the adaptation of cancer cells to mechanical stress, their response under confinement has been extensively studied. Several groups identified an alteration of cell mitosis under such confined situations (11, 12), with modified cell cycle progression and division (13), the presence of multi-daughter events, daughter cells of unequal size, and induction of cell death (14). Moreover, such flattening was shown to promote the G1-S transition (15), as well as the G2-M transition (16), thus fostering cell cycle progression. Under such confined conditions, the nucleus was highlighted as a mechanosensor (17–19), triggering a mechanism of nuclear repair (20) and rescue of DNA damage (21). Upon squeezing, the nucleus shortly unfolds its inner lamin envelope, activating the cytosolic phospholipase A2 (cPLA2)-Arachidonic acid (AA) pathway (17), as well as myosin II contractility and switches towards a motile phenotype (18). Nuclear blebs and chromatin herniation can also occur, leading to transient events of nuclear rupture and repair (20, 22). However, all these data focus on short-term cellular response to confinement, and studying cancer cell fate under several days of confinement remains challenging.

In this study, we investigated whether cells survive and adapt to prolonged squeezing. We showed that they do survive thanks to a global nuclear adaptation that takes place in mitosis. We demonstrated that nuclei relax the applied stress at mitosis by decreasing both their nuclear volume and envelope tension. Both the tension sensor cPLA2 and the contractility machinery were required to achieve proper adaptation. All these changes were conserved by the daughter cells of the next generations, establishing a new state of nuclear homeostasis. Together our results reveal a mechano-adaptation mechanism during mitosis, leading to a decrease in nuclear size..

## Results

### Cancer cells adapt and survive different levels of prolonged confinement by altering their nuclear volume

To investigate whether and how cells adapt to prolonged squeezing, we modified our previously published agarose-based confiner (23), based on a single level of confinement, to subject a cell population to multiple levels of confinement simultaneously (**Fig. 1A**). This system enabled us to impose four different levels of uniaxial deformations (with four different heights from 3 to 9 µm, with a central control zone where no deformation was imposed, **Fig. 1B-C**). After 2 and 24 h, cells displayed a constant decrease in cell and nuclear height according to the levels of confinement (**Fig. 1D-E**, see Materials & Methods for the measure principle), corresponding to an imposed initial nuclear deformation along the z-direction (*ɛ*_*Nz*_) from 41 ± 3% to 70 ± 2% for slight to very strong confinement, respectively (**Fig. 1F**).

**Figure 1:**
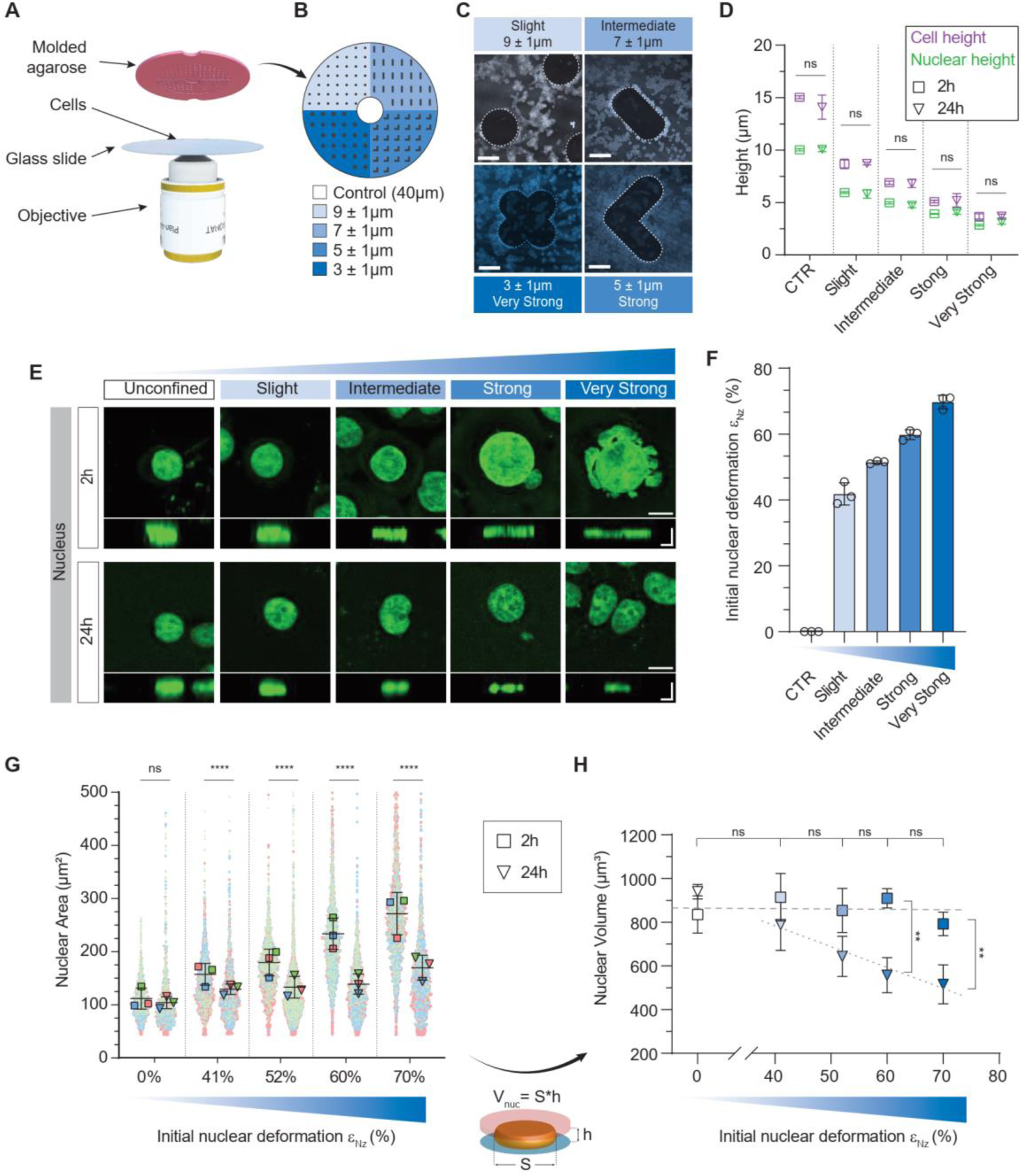
Cancer cells adapt to prolonged confinement by decreasing their nuclear volume. (**A**) 3D representation of our confinement device showing the molded agarose pad. (**B**) Schematic diagram of the multi-height pad with the theoretical heights of each pillar. (**C**) Confocal images of cells and the shaped pillar of each confined zone. *Scale bar = 250 µm*. (**D**) Quantification of cells and nuclear height by z-stack confocal imaging, under each confined zone. *N = 3 experiments with n = 30 cells; Mann-Whitney test*. (**E**) Representative confocal images of nuclei stained with NucGreen in XY (*Scale bar = 10 µm*) and its orthogonal view (scale bar = 5 µm) at 2 h and 24 h under each level of confinement. (**F**) Initial nuclear deformation in percent for each degree of confinement defined as 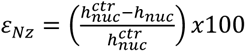. (**G**) SuperPlot quantifying the projected nuclear area. In each condition, the 3 colors represent 3 distinct experiments with their respective individuals and average value represented by small and large dots respectively. *N = 3 experiments with n > 5,000 cells/experiments; Welch’s test. Welch’s ANOVA test was also performed between each confined condition at 24 h and was not significant*. (**H**) Nuclear volume quantification. *N = 3 experiments with n>5,000 cells/experiments; Welch’s test. Ordinary one-way ANOVA was carried out between each confinement condition at 2 h and was not significant*.

We first focus on cell nuclear-projected area and nuclear volume. Different time points were chose. First, 2 h to confirm the lack of nuclear volume adaptation already reported for an imposed deformation of the order of the hour (17, 18), then 24 h, which corresponds to the cell cycle duration of the HT-29 cell line used in this study. We saw that at 2 h of confinement, the nuclear-projected area increased with the level of confinement (**Fig. 1E, G**). The volume of each nucleus, computed by approximating the shape of the nucleus to a cylinder remained constant for all imposed deformations (**Fig. 1H**, squares). Intriguingly, at 24 h of confinement, the nuclear-projected area decreased, resembling that of the control condition (**Fig. 1G**, triangles). This decrease in nuclear-projected area corresponded to a proportional decrease in nuclear volume after 24 h of confinement (**Fig. 1H** triangles), with a loss of nuclear volume from 20 ± 6 % to 34 ± 15 % for slight to very strong confinement, respectively (**Fig. S1A**). Interestingly, this loss in nuclear volume was inversely proportional to nuclear deformation *ɛ*_*Nz*_ (**Fig. 1H**, dotted line). In addition, the distance between the cell membrane and the nuclear envelope was significantly reduced with confinement (**Fig. 1D**, **Fig. S1B**) and accompanied by the re-localization of the contractility machinery (Phosphorylated Myosin Light Chain (p-MLC) staining) from above the nucleus to the side, indicating a cortex rearrangement (**Fig. S1C**). These data support and reinforce our previous findings (23), that cells survive strong squeezing even for a long-term period of 24 h. We provide evidence here that at 24h cells regulate their nuclear volume depending on the level of confinement.

### Nuclear volume adaptation to long-term confinement occurs during mitosis

We then wondered which phase of the cell cycle contributed to this adaptation. It was reported that mechanically confined cells, unable to adopt a rounded shape prior to division, displayed a stressed cell division with delayed mitosis, multi-daughter mitosis events, unevenly sized daughter cells, and induction of cell death (11, 12). Similarly, in our system, we observed a gradual increase in abnormal divisions and mitosis duration with the level of confinement (**Fig. S2A-B, Movie S1**). We further tested if the nuclear volume was conserved throughout mitosis, i.e., if the volume of the sum of the two daughter cells after division was equal to the volume of their mother cell. We focused our analysis on normal division, and only cells capable of progressing through their next cell cycle to mitosis were considered. Intriguingly, nuclei were smaller after the first division under confinement (**Fig. 2A**), with a volume ratio decreasing gradually with increasing nuclear deformation, similar to the one observed at 24 h on fixed samples (**Fig. 1H, 2A**, and **Fig. S2C** for the fit). The loss of volume during mitosis reached up to 60 ± 15% for the strongest confinement (Fig. 2A). The fact that we obtained the same proportionality on fixed (**Fig. 1H**) and live (**Fig. 2A; Fig. S2C**) samples confirmed the robustness of our results. Conversely, during the second division under confinement, we observed no loss of nuclear volume (**Fig. 2B**). Hence, the nuclear volume seemed to reach homeostasis after the first confined division and did not shrink further during a second confined division. Therefore, mitosis appears to be the main checkpoint in the observed long-term adaptation of nuclear volume.

**Figure 2:**
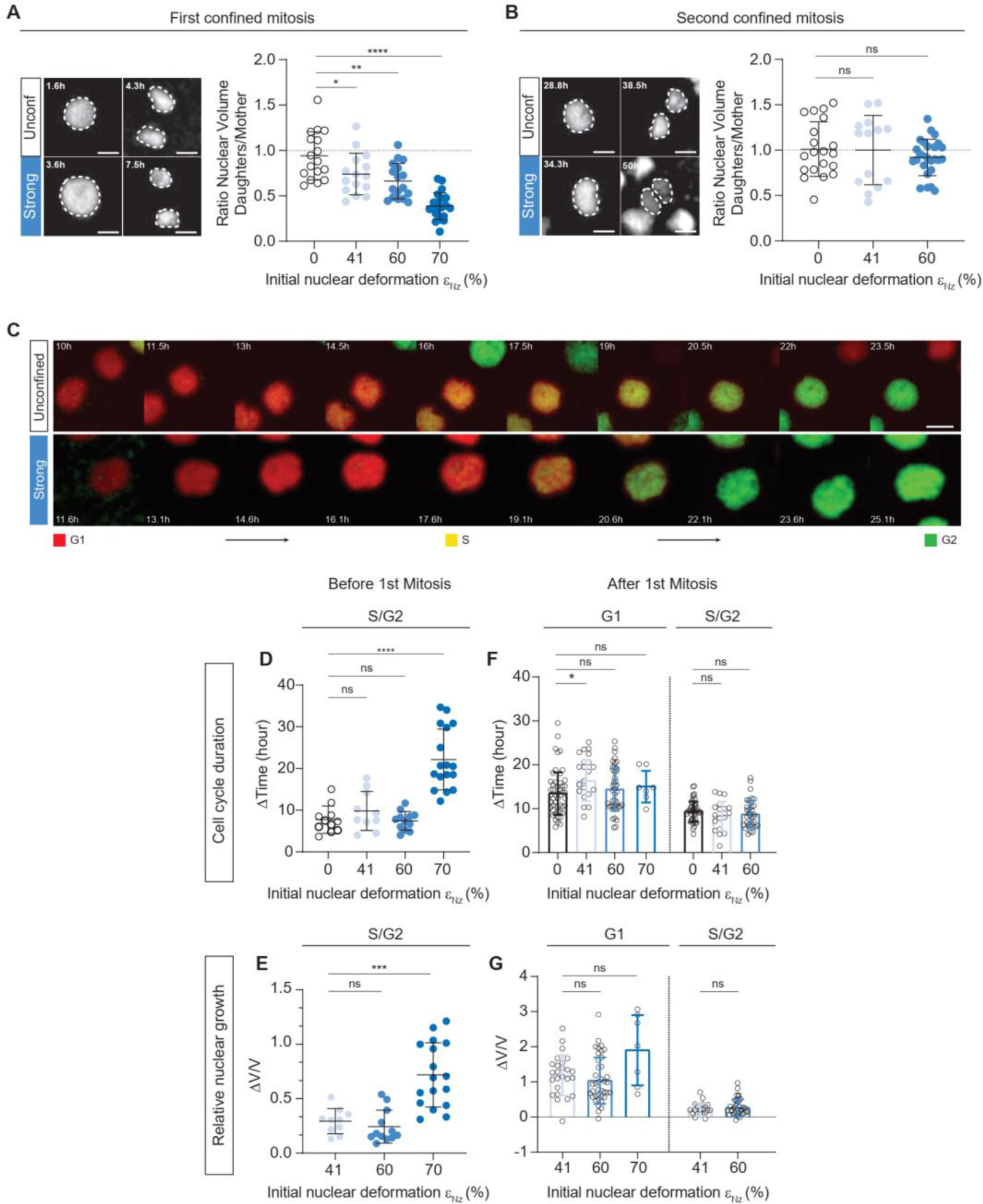
Nuclear volume adaptation occurs during mitosis setting a new state of homeostasis. **(A-B)** Quantification of the size of daughter nuclei over their respective mother, under several levels of confinement during the first confined mitosis **(A)** and the second confined mitosis **(B)** and representative images of nuclei at the end of the G2 phase (just before mitosis) and its respective daughter nuclei just after mitosis (beginning of G1 phase), for control and strong confinement conditions during the first confined mitosis **(A)** and the second confined mitosis **(B)**, scale bar = 10 µm. N = 2 experiments with n > 15 cells per condition, unpaired t-test. **(C)** Time-lapse images of HT-29 nuclei expressing the FUCCI fluorescent cell cycle reporter, without confinement (control) or under strong confinement (60% nuclear deformation). Scale bar = 10 µm. **(D-F)** Quantification of the duration of HT-29 Fucci cells under confinement during the S-G2 before the first division *(D)* and during the G1 and S-G2 phase after the first division **(F).** N = 2 experiments with n = 50 cells **(D)** and n > 200 cells **(F)**, unpaired t test. **(E-G)** Quantification of the nuclear relative growth of HT-29 Fucci cells under confinement during the S-G2 before division **(E)** and during the G1 and S-G2 phase after the first division **(G)**. N = 2 experiments with n = 39 cells, unpaired t test for **(E)** and n > 200 cells, Mann-Whitney test for **(G).**

### Confined mitosis sets a new nuclear state of homeostasis

Next, we investigated whether cell cycle progression was also modified. To this end, we conducted time-lapse experiments over 3 to 4 days on live cells, by confining HT-29 expressing the FUCCI fluorescent cell cycle reporter system (**Fig. 2C**). We initially observed that the nuclear volume at G2 entry was similar for slightly and very strongly confined cells, suggesting that the G1 phase was not or weakly affected in HT-29 cells (**Fig. S2D**). Then, the time spent in G2 before the first division increased only for cells under very strong confinement (**Fig. 2D**) and led to an increase in relative nuclear growth with a constant growth rate during the G2 phase (**Fig. 2E** **and Fig. S2E-F**). This increase in size and growth might be due to an increase in nuclear import following a rise in nuclear envelope tension and uncoupled nuclear and cytoplasmic volumes as proposed by Pennacchio et al. (24–26). Conversely, slighter levels of confinement did not impact cell cycle progression (**Fig. S2E-F**), in accordance with the tension threshold described by Lomakin et al. and Venturini et al. (17, 18).

We then focused on the cell cycle taking place after the first confined division. Surprisingly, for daughter cells, we found that confinement did not significantly change the duration of the whole cell cycle (**Fig. S2I**) and neither that of its subpart (G1, S-G2, **Fig. 2F**). Similarly, relative nuclear growth remained unchanged under confinement during the entire cell cycle (Fig. S2J), and during its subpart (G1, S-G2, Fig. 2G), leading to similar nuclear growth rates regardless of the level of confinement (Fig. S2K), with no significant difference in their subparts (Fig. S2H). These results clearly show that after the first confined division, neither cell cycle progression nor growth rate were affected by confinement, cells thus adapted to confinement during the first confined mitosis and reached a new homeostatic state.

### Mitosis causes the relaxation of the nuclear envelope tension under confinement

As observed, the nuclear volume remains constant for short timescales (2 h), but due to the applied confinement and geometrical considerations, the nuclear envelope (NE) unfolds, and its tension increases (27), which can lead to nuclear protrusions (28–30). Nuclear protrusions are dynamic structures whose membrane can rupture – that can be visualized by leakage of NLS-RFP dye to the cytoplasm (**Fig. S3A**, arrows) – and repair (20). To investigate whether nuclear tension can be regulated under prolonged confinement, we first quantified the nuclear envelope folding from lamin A/C immunostaining (**Fig 3A**) by defining the positive area as the excess area of the lamina envelope compared to the nuclear area (see methods and **Fig. S3B**). In control conditions, the nuclear envelope was folded, reaching similar levels at 2 h and 24 h (56 ± 1.5% and 57 ± 2% respectively, **Fig. 3B**). Under strong confinement (60% nuclear deformation), a complete unfolding of the nuclear envelope was observed at 2 h (**Fig. 3A**), evidenced by a radical decrease in the positive area (**Fig. 3B**, down to 20 ± 1%). At 24 h, a refolding of lamin A/C was observed (**Fig. 3A**), with an increase in the positive area (34 ± 1%, **Fig. 3B**), suggesting that nuclei have adapted to lower their nuclear tension.

**Figure 3:**
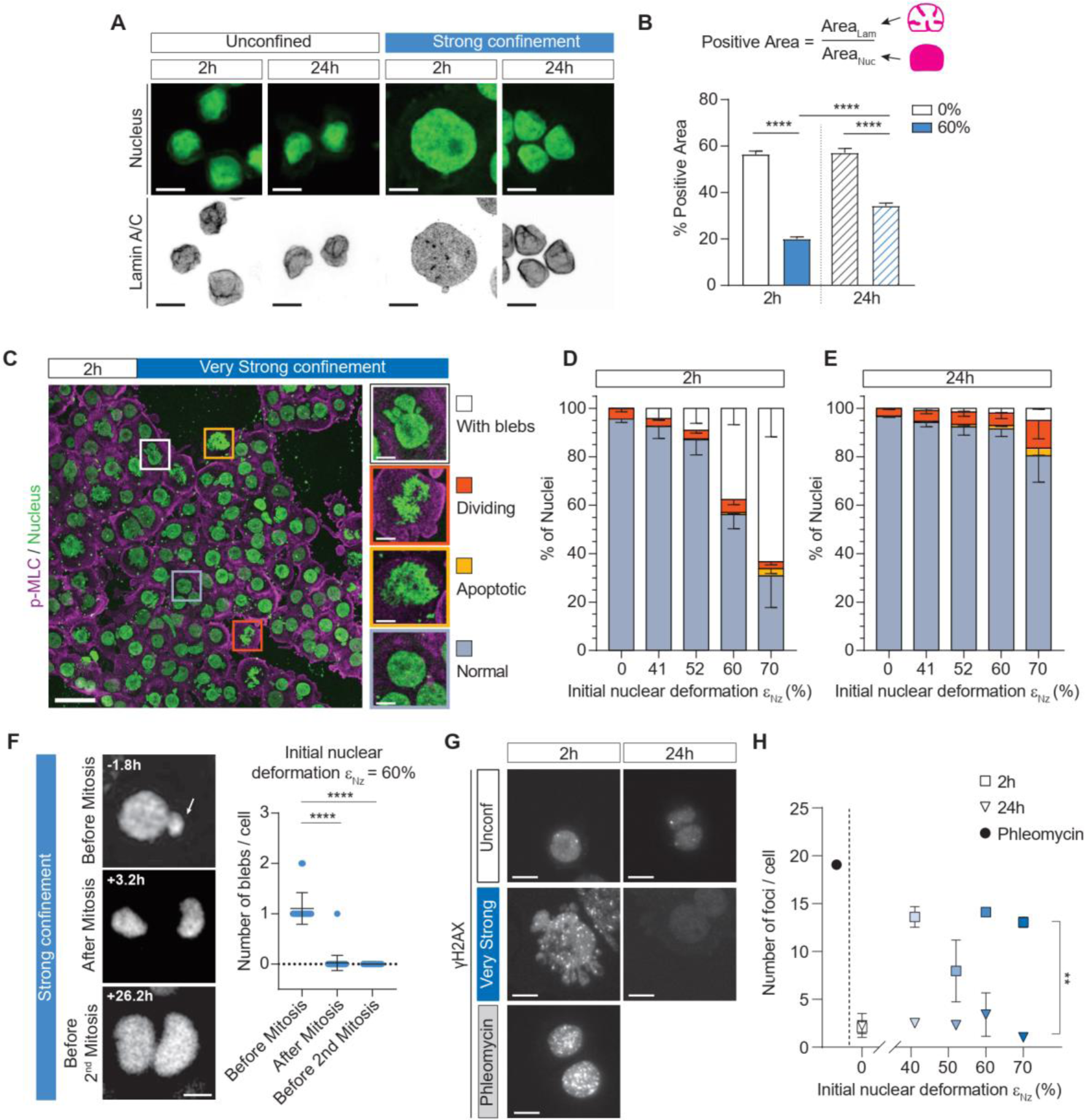
Mitosis regulates nuclear tension under prolonged confinement. **(A)** Representative confocal images (max intensity projection) of stained nuclei and lamin A/C without confinement (control) or under 60% nuclear deformation (strong confinement) at 2 h and 24 h. Scale bar = 10 µm. **(B)** Quantification of the positive area of the nuclear envelope (lamin A/C) which represents the folded nuclear envelope. N = 3 experiments with n > 30cells/experiments; mean ± SEM; Mann-Whitney test. **(C)** Max projection confocal images of stained nuclei and pMLC at 2 h under strong confinement, scale bar = 50 µm. Cropped nuclei correspond to nuclei with blebs, dividing and apoptotic cells and normal nuclei, scale bar = 10 µm. **(D, E)** Percentage of HT-29 cells presenting nuclear blebs, dividing, apoptotic cells, or normal nuclei in control condition or under confinement, at 2 h **(D)** and 24 h **(E).** N = 3 experiments with n > 5,000cells/experiments. **(F)** Sequential images of a blebbing nucleus under strong confinement before and after mitosis, NEB is set here as a reference time t = 0 h, scale bar = 10 µm and quantification of the number of blebs before and after mitosis under strong confinement. N = 3 experiments with n = 77 cells, Mann-Whitney test. **(G)** Representative images of nuclei and DNA damage, shown by stained γH2AX foci. Phleomycin is a drug that induces DNA damage used here as a positive control. Scale bar = 10 µm. **(H)** Quantification of the number of foci/cell, representative of the DNA damages according to an increase in nuclear deformation. N = 2 experiments with n > 2000; mean ± SEM; unpaired t-test. Ordinary one-way ANOVA was also performed on all 24 h confinement conditions and was no significant.

As a proxy for nuclear envelope tension, we analyzed the evolution of nuclear protrusions. Under the strongest levels of confinement (60% and 70% nuclear deformation), we observed nuclear protrusions at 2 h (**Fig. 1E**, strong-very strong confinement panel, and **Fig. 3C**). Such protrusions had a limited contribution to the increase in nuclear-projected area, as the increase remained significantly different even if protrusions were dismissed to compute the projected area (**Fig S3C**). The proportion of nuclei displaying protrusions (indiscriminately called blebs in the rest of the manuscript for simplicity), as well as dividing, apoptotic, and normal nuclei was quantified according to their morphology (**Fig. 3C**, insets). Under slight confinement (41% nuclear deformation), blebbing nuclei represented only 4.2 ± 4.7% of all nuclei (**Fig. 3D**). However, the proportion of these nuclei gradually increased with confinement and represented 37.7 ± 6.7% and 63.4 ± 11.8% of all nuclei for 60% and 70% of nuclear deformation, respectively (**Fig. 3D**). Strikingly, at 24 h of confinement, those proportions dropped to less than 10%, and were mirrored by an increase in normal nuclei proportions (**Fig. 3E** **and Fig S3D**). The number of blebs per cell and the size of these blebs were also drastically reduced at 24 h under strong confinement (**Fig. S3E-H**). This change in nuclear phenotype at 24 h is in accordance with the reduction in nuclear volume and the refolding of the nuclear envelope.

As nuclear volume adaptation occurs after mitosis, we investigated whether this event could also play a role in the loss of nuclear blebs. Using time-lapse experiments, we quantified nuclear blebs before and after confined divisions. Before mitosis, the mean number of blebs per cell was 1 ± 0.3 for cells under strong confinement (**Fig. 3F**, 60% nuclear deformation), and reached 3 ± 2.2 for very strong confinement (**Fig. S3I**, 70% nuclear deformation). Surprisingly, immediately after mitosis, the mean number of blebs per cell was close to 0 for both strong and very strong confinement (**Fig. 3F, Fig S3I, Movie S2**). In addition, we observed no blebs before the second mitosis (**Fig. 3F**). Loss of nuclear blebs is clearly linked to mitosis, suggesting that nuclear volume and nuclear envelope tension are tightly coupled, and supports the hypothesis that mitosis is a key regulator of nuclear envelope tension.

As the presence of nuclear blebs is also associated with DNA damage, we quantified such damages using γH2AX immunostaining. At 2 h of confinement, DNA damage appeared, as evidenced by an increase in the number of γH2AX foci per cell (marker of double-strand breaks, **Fig. 3G**). This confinement-induced DNA damage was repaired at 24 h, with a number of foci per cell in confined conditions back to basal levels (**Fig. 3H**). The mechanism(s) behind such repair is likely coupled to nuclear adaptation, either during cell division itself or due to the relaxation of the nuclear envelope tension at mitosis.

### Cells under confinement regulate their apparent nuclear surface at mitosis

To gain further insights into the mechanism at play, we wondered what physical parameter is involved in the nuclear adaptation described above. We hypothesize that the nuclear volume can be set by the total amount of nuclear envelope or by the apparent surface area of the nucleus (nuclear surface area excluding the membrane ruffles). First, we estimated the total surface area of the nuclear envelope *S*_0_ from the apparent surface area of confined nuclei at 2h, assuming the total unfolding of the nuclear envelope (**Fig. 4A**). Taking S_0_ as constant, we calculated the volume V_0_ a nucleus with a tensed nuclear envelope would have for various nuclear deformation (Total available surface model, **Fig. 4B**, pink line). Second, we estimated the apparent surface area S_app_ of unconfined nuclei using simple geometric assumptions (see supplementary text and **Fig. 4A**). Assuming a constant apparent surface area S_app_, we computed the corresponding nucleus volume V_app_ for various degrees of nuclear deformation (Apparent surface model in **Fig. 4B**, green line). We found that the volumes calculated using the apparent surface model closely matched the experimental volumes measured after 24 h under confinement (**Fig. 4B**, points). This validates the second hypothesis and shows that the apparent surface area is a crucial parameter for nuclear volume adaptation.

**Figure 4:**
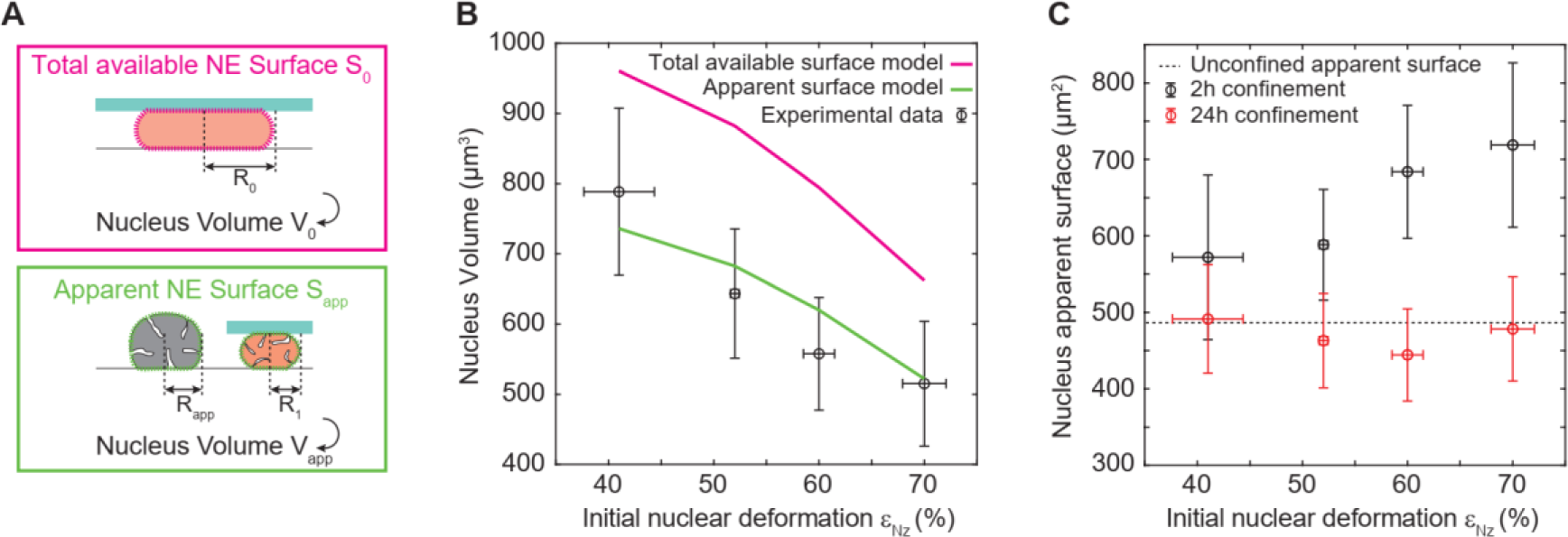
A new state of nuclear homeostasis is determined by the apparent nuclear surface at mitosis exit. (**A**) Schematic representation of the geometric model, showing the apparent nuclear envelope surface area S_app_ and the unfolded nuclear envelope surface area S_0_ (**B**) Calculations of nuclear volume using the apparent surface area (green) and the total surface area (pink). The total surface area is obtained from the tensed nuclear envelope measured under intermediate confinement at 2 h. Experimental values are shown in black. (**C**) Nuclear apparent surface area as a function of the initial nuclear deformation *ɛ*_*Nz*_ at 2 h and 24 h under confinement. The dashed line shows the apparent surface measured for unconfined cells.

To further validate this hypothesis, we also computed the apparent surface area S_app_ from the experimental z-stacked images (assuming a spherical or cylindrical geometry for the unconfined and confined state, respectively). The calculated apparent surface area of confined nuclei *S*_*app*_, passively increased after 2 h under confinement (**Fig. 4C**, black points), while it reached a constant value at 24 h (**Fig. 4C**, red points), very close to the constant apparent surface area computed for unconfined cells (**Fig. 4C**, dotted line). Together, these results suggest that the new homeostatic volume reached by cells after mitosis might be set by the apparent surface area of the nuclear envelope and advocates for a role of nuclear tension sensing that would govern such an apparent surface area.

### Nuclear adaptation to confinement requires the tension sensor cPLA2, but also actomyosin contractility

We then further investigate the molecular actors involved in this adaptation. As the key role of cPLA2 in the sensing of nuclear envelope tension has been highlighted in different contexts (17, 31, 32), we tested whether nuclear volume adaptation was preserved upon cPLA2 inhibition (with AACOCF3). In addition, as cPLA2 is also involved in the regulation of myosin II contractility in a calcium-dependent manner (17, 18), the nuclear volume adaption was also tested upon depletion of intracellular calcium (with BAPTA-AM and 2APB) or inhibition of myosin II (with Blebbistatin). Upon inhibition of cPLA2, myosin II, or upon depletion of intracellular calcium, no further decrease in volume at 24 h was observed (**Fig. 5A**).

**Figure 5:**
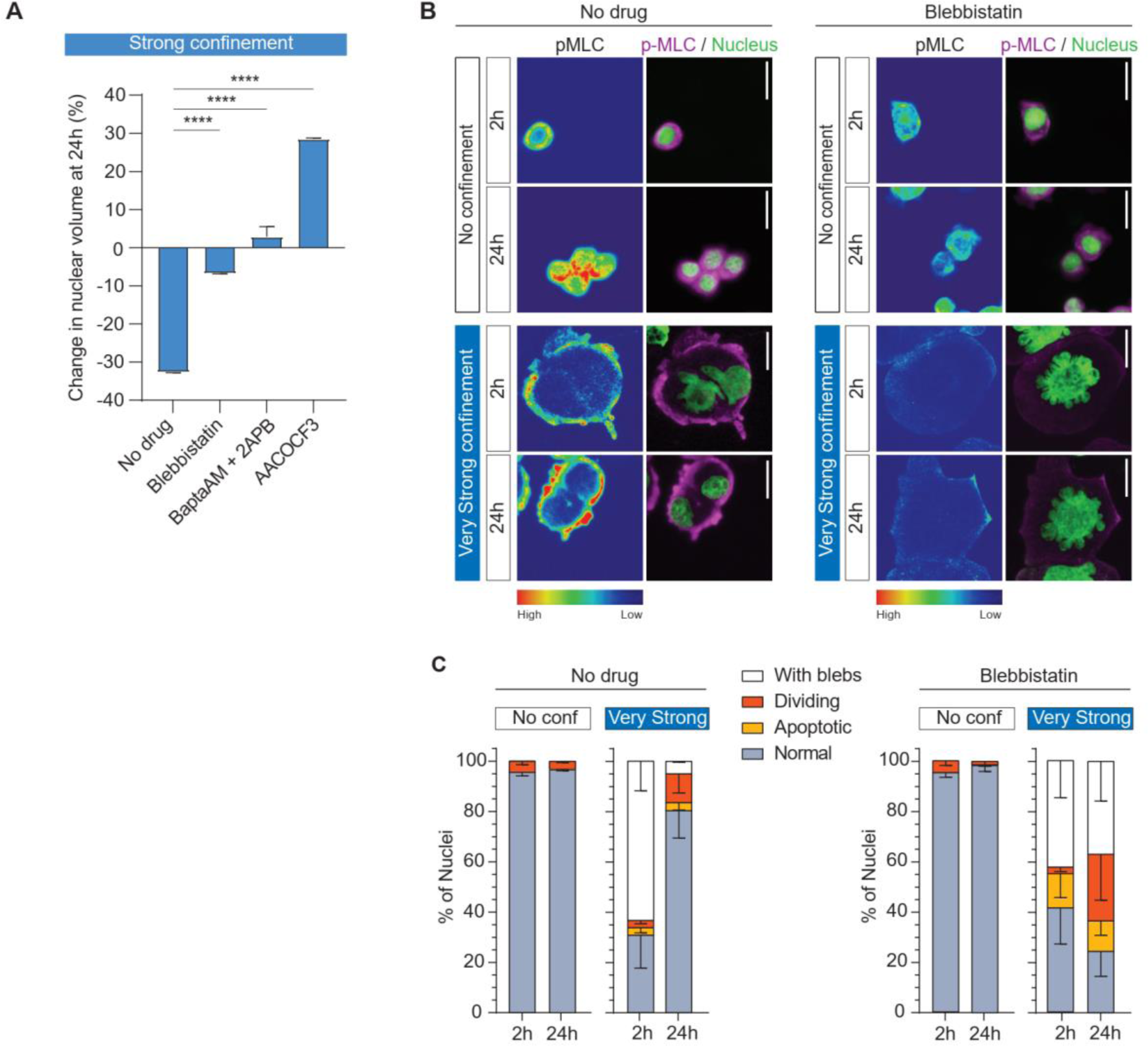
The long-term nuclear adaptation is defective with altered actomyosin cortex. (**A**) Quantification of the nuclear volume changes (in percentage) between 2 h and 24 h without confinement and under strong confinement with several drugs. *N = 3 experiments with n>2,500 cells, mean ± SEM, Welch’s t-test*. (**B**) Representative images of stained nuclei and phospho-Myosin Light Chain either with or without Blebbistatin at 2 h and 24 h, without confinement or under very strong confinement. *Scale bar = 20 µm*. (**C**) Quantification of nuclei with blebs, dividing, apoptotic, or normal without confinement or under very strong confinement at 2 h and 24 h, with no drug or with Blebbistatin. *N = 3 experiments with n = 9,818 cells*.

From this small pharmacological screen, we therefore concluded that the nuclear adaptation observed during mitosis requires nuclear tension sensing through cPLA2 and activation of actomyosin contractility. The latter is a well-known transducer of stress to the nucleus, inducing deformation and blebbing (28). To further confirm that actomyosin contractility was mandatory for nuclear envelop tension relaxation, we quantified the number of blebs in the presence of Blebbistatin. We observed that a large number of blebs were still present after 24 h of confinement (**Fig. 5B-C**). Similarly, inhibiting contractility before applying the confinement (Blebbistatin added 1 h before confinement) also prevented the regulation of nuclear volume and blebs (**Fig S4 A-D**). These results highlight the interplay between contractility, nuclear tension, and nuclear volume regulation in setting nuclear homeostasis upon prolonged confinement.

## Discussion

Deciphering how the nucleus reacts to long-term confinement is of primary importance in the context of cancer progression and response to treatment (33, 34). Using an imposed deformation, we show that the nucleus regulates both its volume and its structure (blebs, lamin A/C folding) within 24 h. We evidenced that this regulation is mediated by mitosis. Nuclei then reach a new state of homeostasis to alleviate nuclear tension, with no further adaptation observed in the next generations. This regulation is an active process requiring the tension sensor cPLA2, intracellular calcium, as well as the contractility machinery.

### Mitosis plays a key role in nuclear adaptation to confinement and alleviation of nuclear envelope tension

The key step in nuclear adaptation to confinement is cell mitosis. We clearly show that the regulation of nuclear volume observed at 24 h of confinement largely originated from volumetric changes at division exit. The duration of confined mitosis can be prolonged due to defects in spindle assembly (12) or leads to asymmetric or multi-daughter division (11), but so far such shrinking of the nucleus during division has never been observed to our knowledge. More strikingly, no further shrinking was observed for the second division under confinement. This important observation suggests a true adaptation to confinement; once adapted, there is no further volume decrease. Intriguingly, the nuclear volume loss reached for the strongest confinement (∼60 % of original nuclear volume) is similar to the range of chromatin volume mentioned in (35). It will be of interest to investigate in future studies whether the volume loss can be attributed to the nucleoplasm volume, so that the volume reached corresponds to the immobile chromatin fraction only. For now, the mechanisms involved remain elusive. Using a mechanistic approach, we showed that the apparent nuclear surface area is the parameter conserved over a long time and therefore during cell division. By regulating the apparent surface area of their nucleus, cells can adjust their nuclear envelope tension to a homeostatic level. Indeed, mitosis also plays a pivotal role in relaxing the nuclear envelope tension generated by confinement as nuclear blebs disappeared completely after the first mitosis, while nuclear envelope folds reappeared in confined conditions at 24 h.

### The critical role of cPLA2 and actomyosin contractility in the regulation of lamina tension and nuclear volume

The nuclear adaptation we evidenced was lost by inhibiting the nuclear envelope tension sensor cPLA2, or by blocking intracellular calcium or contractility. These results strongly highlight the pivotal role of nuclear tension in nuclear homeostasis. Of note, so far cPLA2 was shown to be activated above a tension threshold (17), whereas in our study, we observed a gradual nuclear adaptation, suggesting that other parameters are used as a confinement readout by the cells in mitosis.

These results are also in accordance with the fact that abnormal actomyosin contractility is associated with changes in cell division, such as defects in spindle positioning (12, 36), failed mitosis, and the appearance of polynucleated cells (11, 12). During cell division, actomyosin contractility participates in cleavage furrow formation (37). A recent study pointed out that myosin-II has indeed a role of mechano-protection, with increase of abnormal mitosis and genetic changes upon myosin-II suppression (38). We report here for the first time a role of myosin-II also in mechano-adapatation of nuclear volume and tension: contractility is required to set nuclear volume and nuclear envelope folding, mostly during nuclear envelope reformation at the end of mitosis.

### Regulation of nuclear repair mechanism is associated with lamina folding and nuclear volume loss

Consistent with previous studies (39, 40), we also evidenced an increase in DNA-damage at 2 h of confinement. Such an increase in DNA-damage could lead to chromosomal instability and mutations (41, 42). However, in addition to lamina refolding and nuclear volume loss, we also observed a decrease in DNA damage at 24 h of confinement. This is in accordance with the reported role of lamin in the DNA damage repair pathway (43) and a lower nuclear envelope tension associated with bleb disappearance. Such repair mechanisms have also been reported in other confinement situations (21), as well as in response to other stresses like radiotherapy (44). As such a repair mechanism is highly correlated with cell-cycle re-entry (21), this is again consistent with a key role for mitosis in such DNA-damage regulation in response to nuclear deformation.

### Outlooks

Our results call for an in-depth analysis of the molecular pathways at play, as well as for deciphering the consequences that such regulation could have in the context of cancer and resistance to therapies. The nuclear to cytoplasmic volume ratio, which is constant within a given population, is most likely to be impacted by confinement and changes in nuclear envelope tension (24, 45, 46), and might be at play in the regulation we describe herein. The import/export regulated by the nuclear pore complex is also a putative important player (25), with important implication for mecanosensitive transcription factors such as the Yes-associated protein (YAP) (47, 48) or MKL1 (49). Lastly, we identified a clear change in lamin unfolding/refolding during this nuclear adaptation.

The interplay between lamin A/C level and matrix stiffness has been clearly established (50, 51), and is coupled to cell’s spreading and hence nuclear tension (imposed by the stress fibers on the nucleus). To better understand the underlying mechanism involved in the refolding of the lamin A/C after mitosis highlighted in this study, it would be of interest in the future to investigate how the amount of phosphorylated lamins and overall level of lamin A/C is changing in mitotic cells submitted to such uniaxial deformation. In addition, lamins and lamin-associated proteins are known regulators of nuclear size (46) and should therefore be further investigated, especially in the context of cancer. Indeed, abnormal nuclei shape (52), together with alterations in the expression of lamin A and lamin B is a hallmark for the prognosis of many tumors (33), including colorectal cancer (53). Low lamin A/C is associated with a decrease in stiffness, irregular shape, and larger nuclei deformation (54) as well as increased metastatic capacities (55). Of note, the HT-29 cell line used in this study presents the p53 mutation (56), has no lamin B2, and low lamin B1 level (57). Loss of lamin B1 leads to nuclear blebs, while it has been reported that nuclear envelope rupture is dependent on p53 (58). The role of lamins and p53 mutations could be addressed by analyzing the nuclear adaptation in other cancer cell lines of different origins, different grades, and more specifically with or without p53 and different levels of lamins expression. The regulation of nuclear size identified in this study could have important consequences on resistance to classical chemotherapeutic treatments that target proliferation. Overall, our findings could pave the way for novel therapeutic strategies against tumor cells.

## Materials and Methods

### Cell culture

HT-29 wild type colorectal adenocarcinoma cells were purchased from the ATCC (HT29 HTB-38). A stable cell line expressing NLS-mKate2 was established using the lentiviral vector IncuCyte® nuclight orange (λ_ex_= 588 nm, λ_em_= 633 nm): the cells were selected using puromycin according to the manufacturer’s protocol. The stable HT29 expressing Fucci (Cdt1-RFP and hGeminin-GFP) cell line was a gift from Dr. Toufic Renno’s team (C. Chaveroux team, CRCL, Lyon, France).

All cells were cultured in Dulbecco’s modified Eagle’s medium (DMEM - Glutamax, Gibco^TM^) supplemented with 10% heat-inactivated fetal bovine serum (FBS; PanBiotech™), 100 units/100µg penicillin/streptomycin (Gibco^TM^). Routinely, all HT-29 cell lines were grown in T-25 cell culture flasks and kept at 37°C and under 5% CO_2_. Culture medium was changed regularly, and cell passage was carried out at 70% confluency. Cell lines were tested for Mycoplasma every 6 months and the tests were negative.

### Drug treatments

The following pharmacological inhibitors and chemical compounds were used: Blebbistatin (Myosin II inhibitor Tocris™) at a final concentration of 100 µM; BAPTA-AM (abcam ab120503n) and 2 APB (intracellular calcium depletion, abcam ab120124), both at a final concentration of 10 µM; and AACOCF3 (cPLA2 inhibitor, abcam ab120916) at a final concentration of 20 µM.

To inhibit cPLA2 or intracellular calcium, agarose molds were incubated overnight with a medium with 20 µM AACOCF3 or with 10 µM BAPTA-AM and 10 µM 2 APB, respectively. Immediately after confinement, the medium was replaced by 500 µL of medium with the corresponding drug(s).

To inhibit cell contractility before confinement, we added 500µL of medium with 100 µM Blebbistatin 1 h prior to confinement.

To inhibit the cell contractility after confinement, 500 µL of medium solution with 200 µM of Blebbistatin was added following confinement (to reach a final concentration within the confiner of 100 µM). Initial rinsing after confinement was also made with a medium containing Blebbistatin at 100 µM.

### Multi-height micro-milled mold fabrication

The height of the micropillars (from 9 µm to 3 µm) determines the level of spatial confinement of cells. The array presents regularly distributed pillars (distance between each pillar of 800 µm) with specific shapes according to their height: the “o” shape for 9 µm; the “-” shape for 7µm; the “L” shape for 5 µm; the “+” shape for 3µm. The field of pillars is surrounded by a solid band of 3.5 mm to stabilize the structure, and two external notches to enable the proper positioning during molding. The pillars were drawn using Autodesk software and the mold was created using the CNC Mini-Mill/GX on brass.

### Agarose molding

Based on the procedure described in Prunet et al.(23), molding was adapted to obtain accurate agarose replicates from the newly developed multi-height brass mold. We obtained a user-friendly, robust, and time-saving system to visualize and analyze the long-term effect of four levels of cell confinement on cancer cell morphology and biology.

First, a solution of agarose diluted in distilled water (2% (w/v)) was prepared by autoclaving at 120°C for 10 min, or 30 s microwave (1000 W). Of note a 2% agarose pad corresponds to a stiffness of 150 kPa (23); in this stiffness range, cells cannot deform the confining agarose gel. All the following steps were performed under a sterile culture hood, and the polycarbonate (PC) holder was UV sterilized before use (20 min on each side under 24 W, 365 nm). 400 µL of the prepared agarose solution was deposited in the prewarmed PC holder (placed on a hot plate at 75°C) and was left at room temperature for 5 min to let the gel set. Then, 500 µL of the prepared agarose solution was deposited on the prewarmed micro-milled brass mold (on the same hot plate temperature) with the PC holder and placed under the press (Fig. S5). The press generates a uniform pressure and ensures the contact between the PC holder and the mold, to ensure proper molding and resolute pillars. After 30 min under press at room temperature, the PC holder was gently removed from the brass mold. Evenly distributed holes were drilled into the gel using a 20G and 16G puncher and through holes present on the PC holder. In total, five 16G holes and four 20G holes were done. The molded agarose gel was then placed in sterile PBS and sterilized under UV light (20 min on each side). The molded agarose can be stored in its PC holder at 4°C in PBS until further use. It was replaced either by culture medium or by drug treatment and incubated at 37°C at least 12 h before the confinement experiment.

### Microfabrication-based confinement of cell populations

Cell confinement was performed using the agarose-based soft cell confiner methods previously described (23) and is now available as Agarsqueezer (https://www.idylle-labs.com/agarsqueezer-by-softconfiner). Briefly, after mounting and autoclaving of the confiner holder with a clean glass coverslip, coverslips were coated with fibronectin 50 µg/mL (Sigma-Aldrich™) (1 h incubation in the system at room temperature and washed 3 times with PBS). Cells were then seeded in the systems overnight. An intermediate cell density level was chosen for all experiments (500 µL of a cell solution at 180,000 cells/mL, corresponding to ∼ 45,000 cells/cm², similar to the range used for cell culture). After cell adhesion, the seeded cells were gently washed three times to replace the medium with pre-warmed fresh culture medium (500 μL). The PC holder containing the molded agarose gel was then placed in the system on top of the cells, and a clamping washer was tightened with a specific clamping tool. The molded gel was then tightly in contact with the glass coverslip supporting the cells. A reservoir of 500 μL of culture medium was added above the plastic holder and the molded agarose was then washed three times (5 min each) with pre-warmed culture medium to get rid of any paracrine factors secreted by the dead cell population situated under the pillars.

### Immunostaining

During confinement, cells were fixed *in situ* with paraformaldehyde (PFA): the cell culture medium was removed, and cells were washed twice with PBS. 4% PFA (Electron Microscopy Sciences™) was added and incubated for 40 min at RT. After incubation, samples were rinsed with PBS - 3% BSA (Sigma-Aldrich™) (3 x 20 min). The agarose gel was then removed and cells were permeabilized using 0.5% Triton X-100 (Acros Organics™) for 10 min at RT. After blocking with PBS - 3% BSA (3×5min) to inhibit non-specific binding of antibodies, cells were incubated overnight at 4°C with primary antibodies diluted in PBS - 2% BSA - 0.1% Triton X-100. After 3 x 5 min washes with PBS, samples were incubated with secondary antibodies for 2 h at RT. The primary antibodies used were: p-MLC (Cell signaling™ at 1/50); LaminA/C (Santa Cruz Biotechnology™ at 1/100), γH2Ax (Santa Cruz Biotechnology™ at 1/200). NucGreen™ Dead 488 (Invitrogen™, 1 drop per 3 ml in PBS); Alexa 647 Phalloidin (Invitrogen™ at 1/500); Alexa 555 Phalloidin (Invitrogen™ at 1/500) were also used. The secondary antibodies used were: Anti-Mouse 647 (Invitrogen™ at 1/500); anti-rabbit 647 (Invitrogen™ at 1/500), anti-mouse 555 (Invitrogen™ at 1/500). Coverslips were then mounted using 2 drops of Fluoroshield™ (Sigma-Aldrich™) and varnish-sealed after overnight hardening at room temperature.

### Confocal fluorescence microscopy

Fixed samples were visualized using a Leica SP5 confocal microscope with a 20x Dry objective (NA = 0.7), a 40x oil immersion (NA = 1.25), a 40x Dry (NA = 0.95), or a 63x oil immersion (NA = 1.25) objectives. Images were collected in sequential mode using averaging at a resolution of 1024×1024. Z-stacks of cells were acquired to precisely measure the height of confinement and nuclear volumes (dz = 0.2 µm or 0.5 µm for each stack).

### Epifluorescence imaging

Fixed samples were also visualized using an inverted microscope Leica DMI8 using epifluorescence microscopy with a 40x dry objective (NA = 0.55).

### Live-cell imaging

Cells were observed with an inverted microscope Leica DMI8 using epifluorescence imaging. Timelapse imaging was performed for 2 to 5 days in a controlled environment (CO_2_, temperature, and humidity), using a 20x dry objective (NA = 0.7). A motorized x-y stage enabled the concomitant recording of up to 20 regions for each system every 10 min.

### Quantitative image analysis

#### Cell height, nuclear area, and volume

were assessed with a homemade routine workflow using both ImageJ/Fiji and MATLAB® software. First, after applying a gaussian blur on nuclei-stained images, nuclei were automatically detected either using the stardist Fiji plugin (59) or using the Otsu’s threshold detection directly in Matlab®. Mask and labeled images obtained were exported in Matlab® to perform automated morphological analyses on individual pre-labeled nuclei. Orthogonal views of both nuclei and phalloidin staining were used to obtain nuclear (ℎ_*nuc*_) and cell heights (ℎ_*cell*_ ), respectively. To do so, we took the full width half maxima of the z-plot profile of ImageJ/Fiji of each orthogonal view.

For the control conditions, the nuclear height is determined using the same method. For the volume however, it was necessary to estimate the volume for the reconstituted volume generated from confocal z-stack, by combining 2DStarDist plugin with a home-made python routine.

For the nuclear area, at least four different areas were imaged and analyzed per condition.

#### Proportion analysis of nuclear blebs

both in live and fixed experiments was conducted using the cell counter plugin of ImageJ/Fiji. To further characterize the nuclear blebbing phenotype, labeled images of nuclei and their blebs were obtained using the stardist Fiji plugin (59) (blebs are automatically extracted from the main nuclei using stardist) We filtered the core nucleus and their blebs based on their respective area. If the area of an object was below 100 µm² for confined conditions, it was considered to be a bleb.

#### DNA damage analysis

was automatically performed using the Find Maxima tool within each nucleus in ImageJ/Fiji.

#### Folding of the nuclear envelope

The percentage of **positive nuclear envelope area** was used as a semi-quantitative parameter as a proxy for the folding of the nuclear envelope (17, 60). It corresponds to the excess area of the lamina envelope compared to the nuclear area (Fig. S3B). The nuclear envelope-positive area was assessed by thresholding the maximum intensity projection images of lamin A/C staining obtained by confocal microscopy (63x objective NA 1.25, Leica SP5) and merging them with nuclei masks. This allowed us to study the laminas within the nucleus and provided information on the proportion of area with laminas within the nucleus (positive area).

#### Live movies

were analyzed manually by using either MTrackJ, or manual tracking and ROI manager tool in ImageJ/Fiji plugin. Volume was calculated using the area of each cell multiplied by the theoretical confinement height measured previously for each confinement zone (Figure 1D). Thus, we could measure nuclear areas over time in each phase of the cell cycle and the corresponding duration of those phases.

For the analysis of volume changes during mitosis, only normal mitosis were considered. Therefore, only cells that were able to progress in their next cell cycle up to another mitosis were taken into account.

For each experiment, at least three different areas within each confined zone of the confiner were analyzed.

### Statistical Analysis

Statistical data were expressed as mean ± standard deviation unless mentioned otherwise. Sample size (n) and the number of repetitions (N) are specified in the text of the paper or figure legends. The statistical significance between experimental conditions was determined using a two-tailed unpaired t-test, Welch’s test, Mann-Whitney test, or ANOVA after confirming that the data met appropriate assumptions (normality distribution, homogeneous variance, and independent sampling). All data were analyzed with GraphPad Prism 8.0. (San Diego, CA, USA). **** P < 0.0001, *** P < 0.001, ** P < 0.01, * P < 0.05.

### Data Availability

Data will be shared using Zotero platforms (raw data extracted from image analysis, a set of typical images, as well as matlab code used for image analysis) **[dataset].**

## Acknowledgments

This work was supported by the Institut Convergence Plascan: ANR-17-CONV-0002, the Institut Universitaire de France (IUF) and the ARC for part of M. Mouelhi salary.

The authors would like to acknowledge C. Chaveroux, Marie Piecyk, C. Duret (CRCL) and S. Joly for the transfection and selection of HT-29-FUCCI cell line, R. Fulcrand for the preparation of 5-µm wafer molds, M. Mercury for the microscope holder and the press. G. Jardiné for his help to compute the nuclear volume of control cells, B. Manship for English editing of the manuscript, as well as M. Piel for fruitful discussions of our results and careful reading of the manuscript.

## Notes

### Competing Interest Statement

The authors have declared no competing interest.

### Summary of Updates

The manuscript has been thoroughly rewritten, and new results obtained after different drug treatment has been added

