## Supplementary material for "Mitosis sets nuclear homeostasis of cancer cells under confinement": Fig.S

### Supporting Information Text

#### Geometrical model.

Confined nuclei can be modeled as a cylinder. Thus, their surfaces were calculated using the following equations.

$$2\pi R_1 h + 2\pi {R_1}^2 = S_{app} \qquad \text{(Equation 1)}$$

With $R_1$, the diameter of the nucleus, h, its height, and $S_{app}$, its apparent surface.

$R_1$, the radius of this cylinder can then be extracted:

$${R_1}^2 + R_1 h - \frac{S_{app}}{2\pi} = 0 \qquad \text{(Equation 2)}$$

With a positive solution:

$$R_1 = \frac{-h+\sqrt{\Delta}}{2} \qquad \text{(Equation 3)}$$

To compute the nuclear envelope surface to add during division, we first calculated the surface of the mother cell using equation 1 and then its volume. Using the obtained volume and equation 1 for two cells, we finally computed the surface of the two daughter cells. For non-confined cells, we used a similar approach with spheres rather than cylinders.

**Fig. S1.**

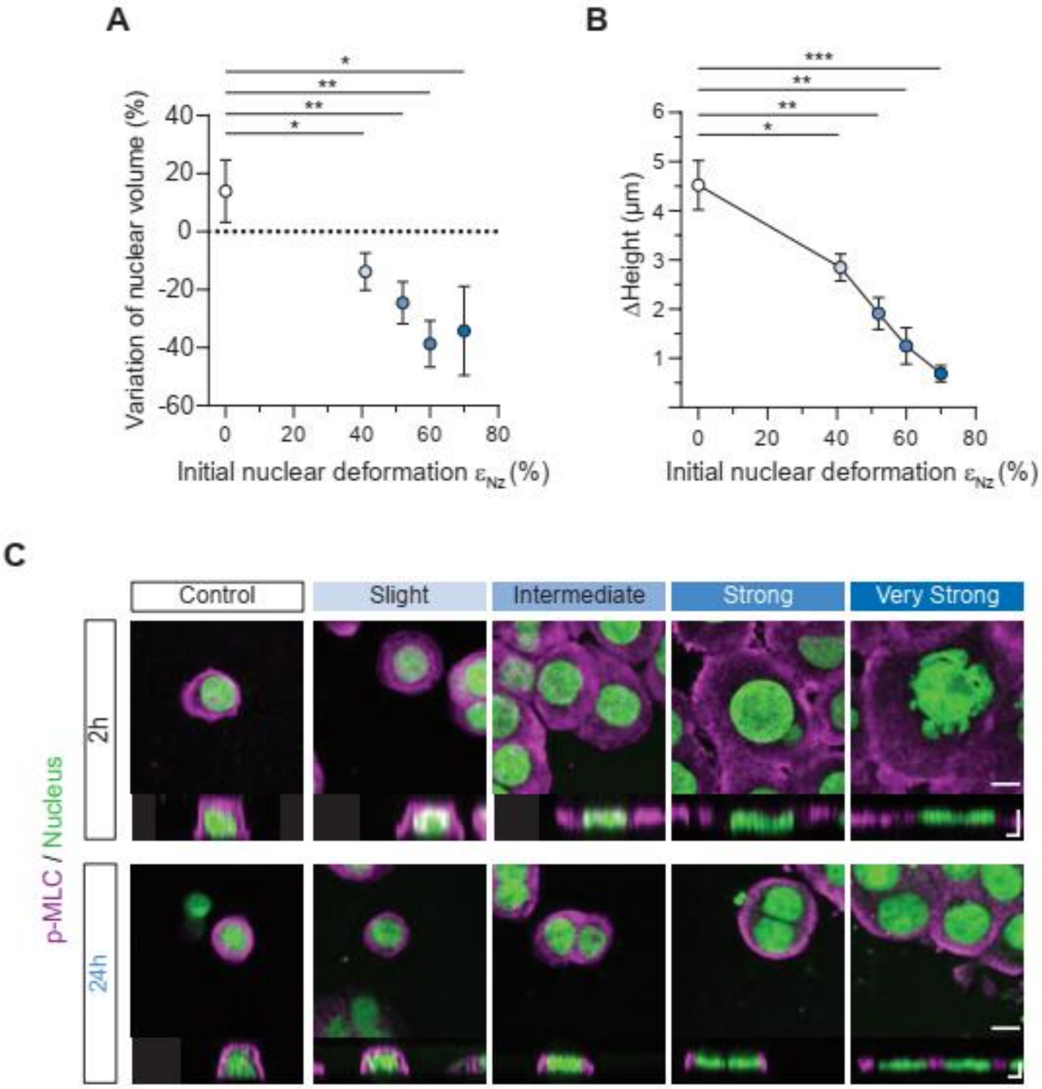


**(A)** Variation of nuclear volume between 2 h and 24 h according to the initial nuclear deformation. *N = 3 experiments with n > 5,000 cells/experiment, unpaired t-test*. **(B)** Difference in height between the nucleus and its cell membrane under increasing initial nuclear deformation. *N = 3 experiments with n > 5,000 cells/experiment.* **(C)** Representative images of stained nuclei and p-MLC in XY (scale bar = 10 µm) and its orthogonal view under increasing confinement (scale bar = 5 µm) at 2 and 24 h of confinement.

**Fig. S2.**

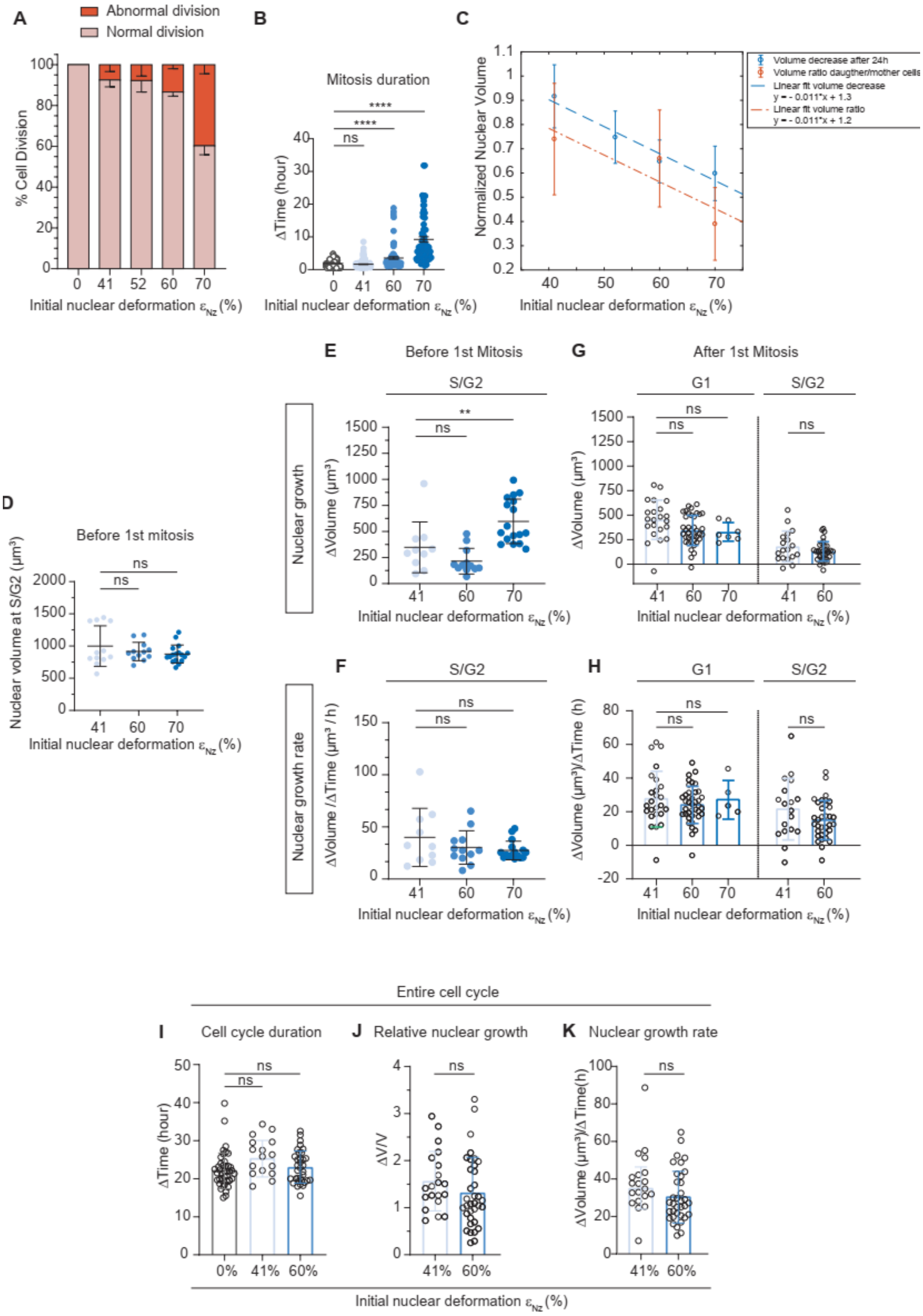
A
Abnormal division
Normal division
% Cell Division
Initial nuclear deformation $\varepsilon_{Nz}$ (%)
B
Mitosis duration
ΔTime (hour)
ns
****
****
Initial nuclear deformation $\varepsilon_{Nz}$ (%)
C
Normalized Nuclear Volume
Volume decrease after 24h
Volume ratio daugther/mother cells
Linear fit volume decrease
y = - 0.011*x + 1.3
Linear fit volume ratio
y = - 0.011*x + 1.2
Initial nuclear deformation $\varepsilon_{Nz}$ (%)
D
Before 1st mitosis
Nuclear volume at S/G2 (μm³)
ns
Initial nuclear deformation $\varepsilon_{Nz}$ (%)
E
Before 1st Mitosis
S/G2
Nuclear growth
ΔVolume (μm³)
ns
**
Initial nuclear deformation $\varepsilon_{Nz}$ (%)
G
After 1st Mitosis
G1
S/G2
ΔVolume (μm³)
ns
Initial nuclear deformation $\varepsilon_{Nz}$ (%)
F
S/G2
Nuclear growth rate
ΔVolume /ΔTime (μm³ / h)
ns
Initial nuclear deformation $\varepsilon_{Nz}$ (%)
H
G1
S/G2
ΔVolume (μm³)/ΔTime (h)
ns
Initial nuclear deformation $\varepsilon_{Nz}$ (%)
Entire cell cycle
I
Cell cycle duration
ΔTime (hour)
ns
J
Relative nuclear growth
ΔV/V
ns
K
Nuclear growth rate
ΔVolume (μm³)/ΔTime(h)
ns
0%
41%
60%
Initial nuclear deformation $\varepsilon_{Nz}$ (%)

**(A)** Percentage of normal or abnormal cell divisions (multi-daughter cells and unevenly sized daughter cells) under increasing confinement. *N = 2 experiments with n = 477 cells*. **(B)** Scatter plot of mitosis duration under confinement. *N = 3 experiments with n = 398 cells; mean ± SEM, Mann-Whitney test.* **(C)** Normalized nuclear volume at 24 h of confinement and nuclear volume after mitosis as a function of initial nuclear deformation and their respective linear fit. **(D)** Using HT-29 cell expressing Fucci, scatter plot of the initial nuclear volume of mother cells at the entrance of the G2 phase, *N = 2 experiments with n = 41 cells, unpaired t-test*. **(E-F)** Using HT-29 cell expressing Fucci, growth **(E)** and nuclear growth rate **(F)** of mother cells nuclear volume during the G2-phase preceding the first mitosis for the different levels of confinement. *N = 2 experiments with n = 39 cells, Mann-Whitney test*. **(G-H)** Using HT-29 cell expressing Fucci, growth **(G)** and nuclear growth rate **(H)** of daughter cells nuclear volume in the G1 and S-G2 phases after the first mitosis for the different levels of confinement. *N = 2 experiments with n > 200 cells, Mann-Whitney test.*
**(I, J, K)** Quantification of the cell cycle duration **(I),** relative nuclear growth **(J)** and nuclear growth rate **(K)** of HT-29 cell expressing Fucci under confinement after the first cell division. *N = 2 experiments with n > 200 cells, unpaired t-test for (I, J) and Mann-Whitney test for (K).*

**Fig. S3.**

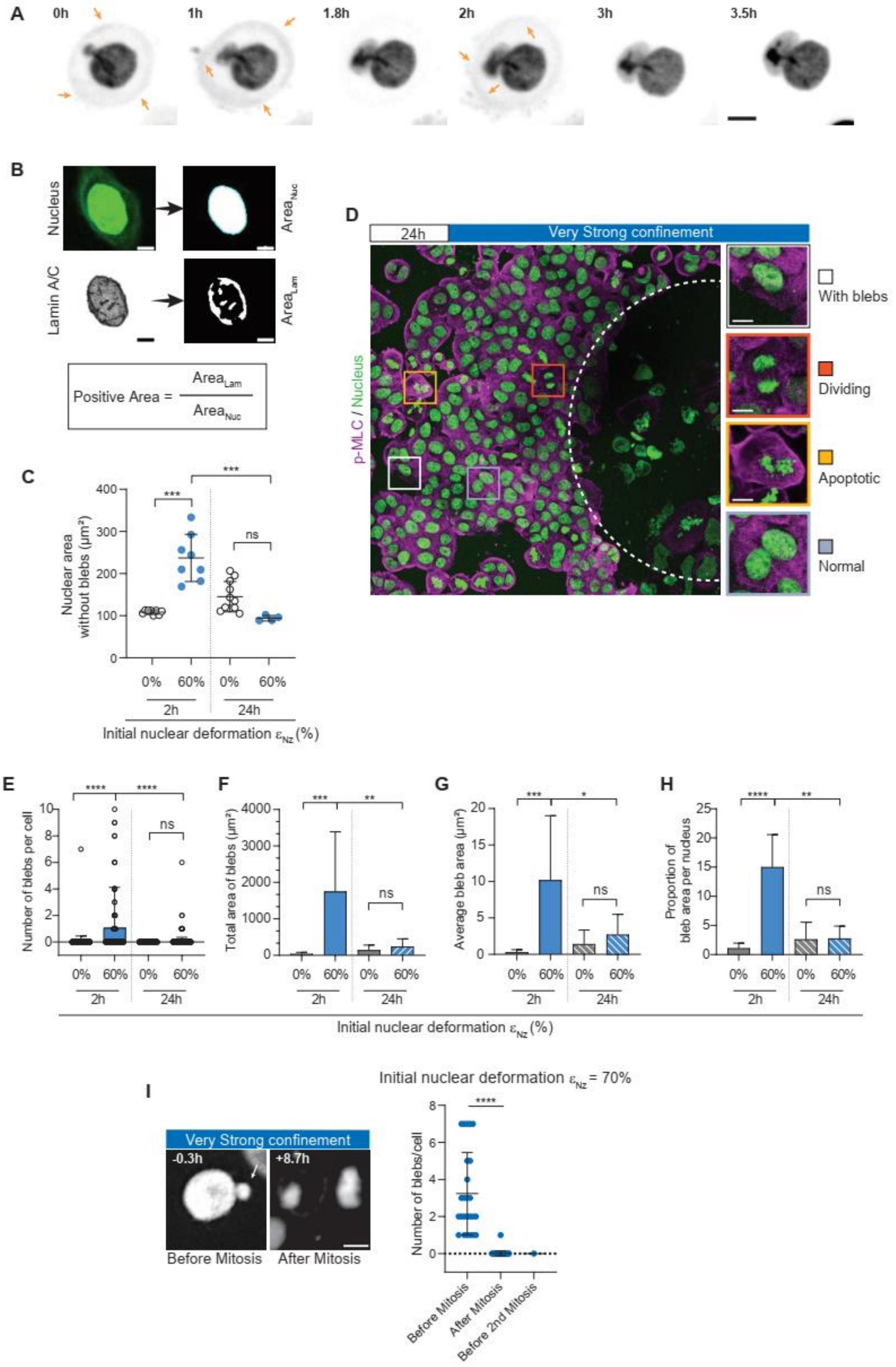
A
0h
1h
1.8h
2h
3h
3.5h
B
Nucleus
Area_Nuc
Lamin A/C
Area_Lam
Positive Area = Area_Lam / Area_Nuc
C
Nuclear area without blebs (µm²)
Initial nuclear deformation ε_Nz (%)
D
24h
Very Strong confinement
p-MLC / Nucleus
With blebs
Dividing
Apoptotic
Normal
E
Number of blebs per cell
F
Total area of blebs (µm²)
G
Average bleb area (µm²)
H
Proportion of bleb area per nucleus
Initial nuclear deformation ε_Nz (%)
I
Very Strong confinement
-0.3h
+8.7h
Before Mitosis
After Mitosis
Initial nuclear deformation ε_Nz = 70%
Number of blebs/cell
Before Mitosis
After Mitosis
Before 2nd Mitosis

**(A)** Sequential images of HT-29 cell expressing NLS-mKate2 under strong confinement. Arrows show NLS-mKAte2 dye leakage within the cytoplasm indicating nuclear membrane rupture. Scale bar = 10 µm. **(B)** Schematic representation of the quantification used of nuclear envelope folding: the positive area of the nuclear envelope (lamin A/C) which represents the folded nuclear envelope. It was assessed by thresholding the maximum intensity projection images of lamin A/C staining obtained by confocal microscopy and merging them with nuclei masks. **(C)** Scatter plot of nuclear area without the blebs at 2 h and 24 h without confinement or under strong confinement. *N = 3 experiments with n = 31 cells, unpaired t-test*. **(D)** Typical Max projection confocal image of stained nuclei and pMLC at 24 h under strong confinement, *scale bar = 50 µm*. Cropped nuclei correspond to nuclei with blebs, dividing and apoptotic cells and normal nuclei, *scale bar = 10 µm*. **(E)** Bar chart of the number of blebs per cell according to the nuclear deformation (0% or 60%) at 2 h or 24 h of confinement. *N = 3 experiments with n = 1,376 cells, unpaired t-test*. **(F)** Quantification of the total area of blebs without confinement or under strong confinement (60% nuclear deformation) between 2 h and 24 h. *N = 3 experiments with n = 36, Mann-Whitney test*. **(G)** Quantification of the average area of blebs without confinement or under high confinement (60% nuclear deformation) between 2 h and 24 h. *N = 3 experiments with n = 36, Mann-Whitney test*. **(H)** Proportion (in percentage) of bleb area per nucleus without confinement or under high confinement (60% nuclear deformation) between 2 h and 24 h. *N = 3 experiments with n = 1,376 cells, Kolmogorov-Smirnov test.* **(I)** Sequential images of a blebbing nucleus under very strong confinement before and after mitosis, NEB is set here as a reference time t = 0 h, *scale bar = 10 µm* and quantification of the number of blebs before and after mitosis under very strong confinement. *N = 3 experiments with n = 90 cells, Mann-Whitney test*.

**Fig. S4.**

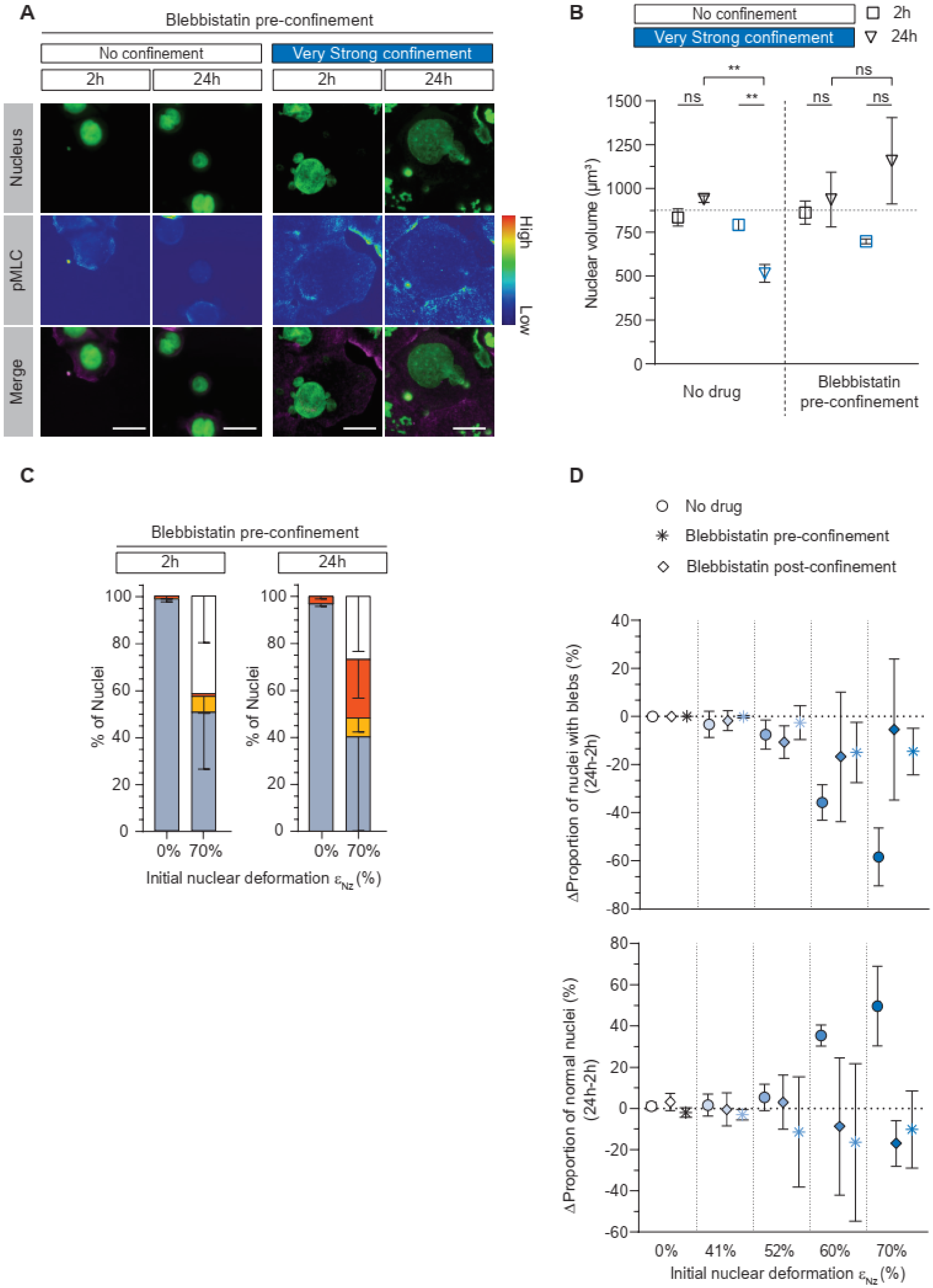
A
Blebbistatin pre-confinement
No confinement
Very Strong confinement
2h
24h
2h
24h
Nucleus
pMLC
Merge
High
Low
B
No confinement
Very Strong confinement
2h
24h
Nuclear volume (µm³)
1500
1250
1000
750
500
250
0
ns
**
**
ns
ns
ns
No drug
Blebbistatin pre-confinement
C
Blebbistatin pre-confinement
2h
24h
% of Nuclei
100
80
60
40
20
0
0% 70%
0% 70%
Initial nuclear deformation ε_Nz (%)
D
No drug
Blebbistatin pre-confinement
Blebbistatin post-confinement
ΔProportion of nuclei with blebs (%) (24h-2h)
40
20
0
-20
-40
-60
-80
ΔProportion of normal nuclei (%) (24h-2h)
80
60
40
20
0
-20
-40
-60
0%
41%
52%
60%
70%
Initial nuclear deformation ε_Nz (%)

**(A)** Representative images of stained nuclei and phospho-Myosin Light Chain (p-MLC) with Blebbistatin – added before confinement – at 2 h and 24 h without confinement or under very strong confinement. Scale bar = 20 µm. **(B)** Quantification of nuclear volume at 2 h and 24 h without drug or with Blebbistatin added before confinement without confinement or under very high confinement. *N = 3 experiments with n = 7,526 cells, mean ± SEM, unpaired t-test (for no drug conditions) or Mann-Whitney test (for Blebbistatin conditions).* **(C)** Quantification of nuclei with blebs, dividing, apoptotic or normal without confinement or under very strong confinement at 2 h and 24 h, with Blebbistatin added before confinement. *N = 3 experiments with n = 7,526 cells.* **(D)** Difference in proportion of normal nuclei and nuclei with blebs between 2 h and 24 h under several conditions: without drug, with Blebbistatin added pre-confinement and with Blebbistatin added post-confinement. *N = 3 experiments with n > 10,000 cells.*

**Fig. S5.**

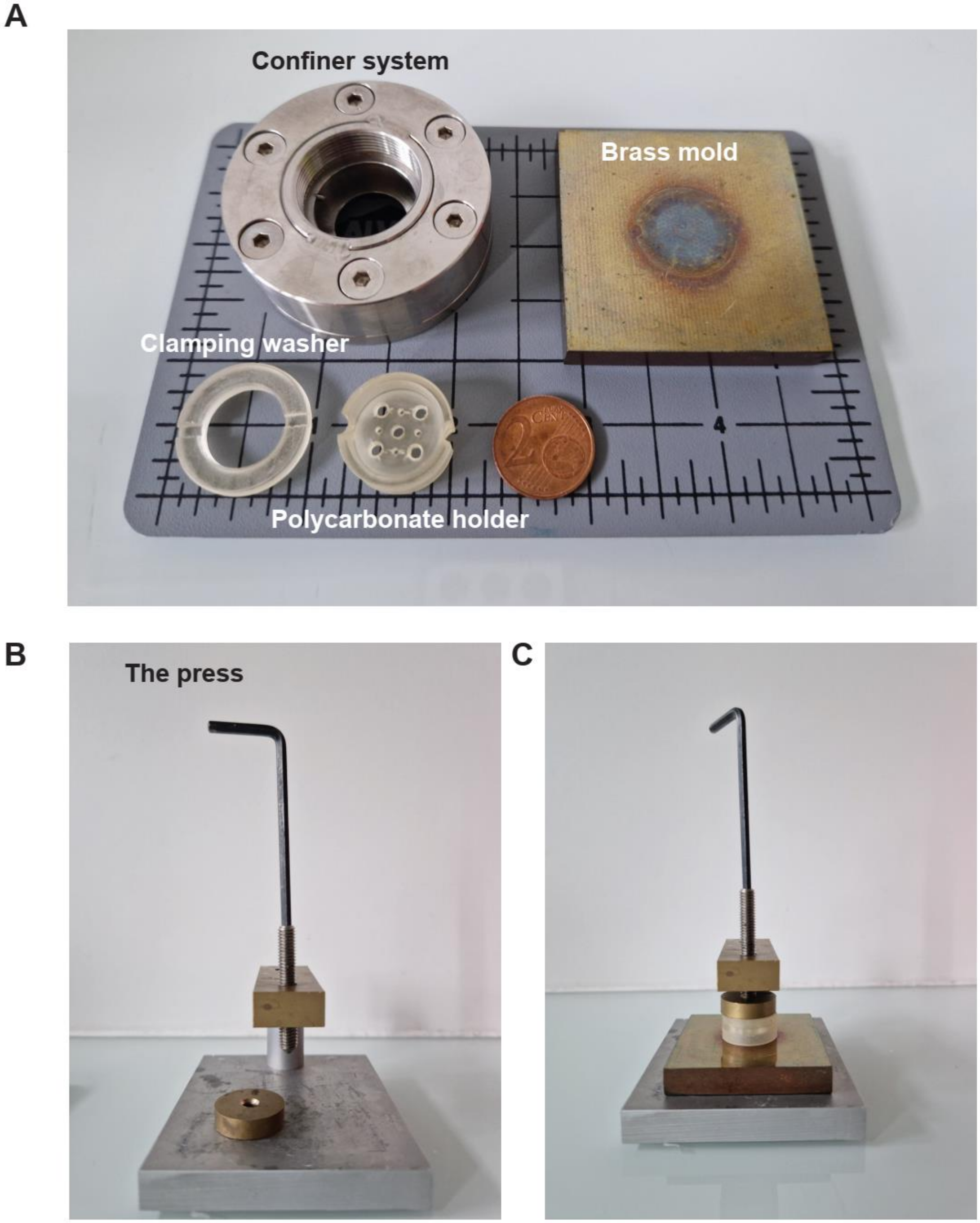


**(A)** Picture of the principal components of the confiner system and their name next to the brass mold. **(B)** Picture of the press used to generate uniform pressure during agarose molding. **(C)** Picture of the assembly process of agarose molding under press.

**Movie S1 (separate file).** Example of abnormal divisions under very strong confinement.

**Movie S2 (separate file).** Example of blebs disappearance after mitosis
